# Lipopolysaccharide administration alters extracellular vesicles in human lung-cancer cells and mice

**DOI:** 10.1101/2020.04.17.046367

**Authors:** Leandra B. Jones, Sanjay Kumar, Courtnee’ R. Bell, Brennetta J. Crenshaw, Mamie T. Coats, Brian Sims, Qiana L. Matthews

## Abstract

Extracellular vesicles (EVs) play a fundamental role in cell and infection biology and have the potential to act as biomarkers for novel diagnostic tools. In this study, we explored the *in vitro* impact of bacterial lipopolysaccharide administration on a cell line that represents a target for bacterial infection in the host. Administration of lipopolysaccharide at varying concentrations to this A549 cell line caused only modest changes in cell death, but EV numbers were significantly changed. After treatment with the highest concentration of lipopolysaccharide, EVs derived from A549 cells packaged significantly less interleukin-6 and lysosomal-associated membrane protein 1. We also examined the impact of lipopolysaccharide administration on exosome biogenesis and cargo composition in BALB/c mice. Serum-isolated EVs from lipopolysaccharide-treated mice showed significantly increased lysosomal-associated membrane protein 1 and toll-like receptor 4 levels compared with EVs from control mice. In summary, this study demonstrated that EV numbers and cargo were altered using these *in vitro* and *in vivo* models of bacterial infection.

## 1. Introduction

Gram-negative bacteria can cause severe illnesses such as pneumonia, meningitis, and bacteremia [1]. These infectious bacteria have become increasingly resistant to antibiotics partly due to their double-membrane structure made from phospholipid and lipopolysaccharide (LPS) [2]. This outer membrane forms a very effective protective barrier, making these bacteria highly resilient to antibiotics [2]. Bacterial LPS is an endotoxin, and a major component of the outer leaflet of gram-negative bacteria that causes inflammatory responses [3]. This barrier protects bacteria from host-immune defenses, mediates direct interactions with both antibiotics and host cell receptors, and initiates events that cause tissue damage in the host [4]. Thus, LPS plays a major role in pathogenesis. The classical LPS molecule is composed of three components: lipid A, an oligosaccharide core (core O), and O antigen polysaccharide [3]. LPS virulence resides in the endotoxicity of lipid A and in the ability of the core O region to provide bacteria with resistance to host immune defenses [3]. Bacterial changes, such as LPS variations, during disease pathogenesis result in immune system response, chronic inflammation, and increased antibiotic resistance [3]. As a result of these variations, molecules (i.e., virulence factors) could potentially be packaged into exported vesicles to promote pathogenesis. These bacterial variations, including modifications to LPS synthesis, are a recurring aspect of infections, regardless of the type of bacteria or the infection site [3]. In general, these changes result in immune system evasion, persistent inflammation, and increased antimicrobial resistance.

Both gram-negative bacteria and their mammalian host cells have the ability to release extracellular vesicles (EVs). Specifically, these EVs are 50–150 nm in size [5], have a density of 1.23–1.16 g/mL, and can transfer molecules (i.e., mRNA, miRNA, RNA, lipids, and proteins) from one cell to another [6, 7]. EVs are found in many biological fluids including semen, breast milk, cerebrospinal fluid, salvia, serum, and urine [5]. Gram-negative bacteria release small EVs from their cell membranes [8], and during infections, these vesicles can package and deliver toxins, as well as other virulence factors, to the host [9]. During infections, EVs can have opposing roles — both initiating an immune response in the host and aiding the spread of the infection through increased pathogenesis [11]. This is due in part to the molecular packaging of EVs being both pathogen-derived and host-derived [7]. EVs isolated from infected cells can function to attenuate, or even eliminate, the spread of infectious disease [10]. These fundamental mechanisms of exosomal molecular packaging and specificity remain unclear and need to be studied further.

Given the importance of EVs both in normal biology and in pathogenesis, they are being studied for their therapeutic potential as biomarkers for disease and disease progression, as immunomodulators, and as molecules for drug delivery. Here, we explored the influence of bacterial LPS (as a gram-negative model) on EVs derived from lung cells cultured *in vitro*, and on EVs derived from an *in vivo* mouse model.

## 2. Materials and Methods

### 2.1 Cell line and culture conditions

A549 lung cells were incubated in Dulbecco’s Modified Eagle Medium F12 media with 10% fetal bovine serum (FBS; Thermo Fisher Scientific, Waltham, MA, USA), 1% penicillin, 1% streptomycin, and 0.5 μg/mL amphotericin in 32 °C with a CO_2_ atmosphere. The cells were grown to 70–80% confluency before proceeding with experiments. For EV experiments, the medium included exosome-depleted FBS. DMEM exosome-free media was prepared using exosome-depleted FBS from System Biosciences (Palo Alto, CA, USA).

### 2.2 Antibodies

Primary antibodies against the following proteins were used: Interleukin (IL) 6 (Developmental Studies Hybridoma (DSHB), Iowa City, IA, USA, 1:1000), Toll-like Receptor (TLR) 4 (Thermo Fisher Scientific, Waltham, MA, USA, 1:2000), TLR7 (Thermo Fisher Scientific, Waltham, MA, USA, 1:2000), Tumor Necrosis Factor (TNF) α (Bioss, Woburn, Massachusetts, USA, 1:2000), HSP90β (Thermo Fisher Scientific, Waltham, MA, USA, 1:1000), CD9 (Thermo Fisher Scientific, 1:1000), CD81 (Thermo Fisher Scientific, Waltham, MA, USA, 1:1000), CD63 (Thermo Fisher Scientific, Waltham, MA, USA, 1:1000), exosome complex exonuclease (RRP44/DIS3) (DSHB, Iowa City, IA, USA, 1:1000), inducible nitric oxide synthase (iNOS) (DSHB, Iowa City, IA, USA, 1:1000) and Lysosomal-Associated Membrane Protein (LAMP) 1 (DSHB, Iowa City, IA, USA, 1:1000). The secondary antibodies used were as follows: HRP-conjugated goat anti-mouse (1:5,000 dilution, Dako Agilent, Santa Clara, CA, USA), HRP-conjugated rabbit anti-hamster (1:2000 dilution, (Thermo Fisher Scientific, Waltham, MA, USA) or HRP-conjugated goat anti-rabbit (1:1,000 dilution, Thermo Fisher Scientific, Waltham, MA, USA).

### 2.3 LPS treatment on A549 cells

Cells were plated at 500,000 cells/flasks and incubated at 24 hours (hrs). After 24 hrs, cells were washed and media was replaced with exosome-depleted media. The experimental flasks were treated with 0.1 μg/mL, 1 μg/mL or 10 μg/mL of LPS for 48 hrs [11]. Following treatment, cell morphology and integrity were visualized and media was collected.

### 2.4 Extracellular vesicle isolation *in vitro*

EVs were harvested from the supernatants of LPS treated cells, A549. The collected media were centrifuged at 300 × g for 10 minutes at 4 °C. The cell culture supernatant was then removed and spun at 2600 × g at 10 mins at 4 °C. Dead cells and cell debris were furthered removed by filtering with either a 0.22 and 0.45-µm filter. 1X PBS was added to the media with 5% sucrose and 1X protease inhibitor and centrifuged at 110,000 × g for 70 minutes (mins) in a SW41Ti swinging bucket rotor at 4°C using a Beckman Coulter Optima ™ L-70K Ultracentrifuge. After 70 mins, the media was decanted and approximately 500 µL of extracellular vesicle pellets were collected and stored at −20 °C. EVs were quantitated using Bradford-Lowry protein quantitation procedure (Bio-Rad Laboratories, Hercules, CA, USA).

### 2.5 MTT cell viability assay

Cell viability was assessed using the 3-(4,5-dimethylthiazo-1-2yl)-2,5-diphenyltetrazolium bromide (MTT) assay (Thermo Fisher Scientific, Waltham, MA, USA). A549 cells were seeded independently in 96-well tissue culture plates (10,000 cells/well) and maintained in culture for 24 hrs prior to treatment. Then, standard media was exchanged for serum-free media and cells were stimulated with LPS (*E. coli* LPS O55:B5, Sigma-Aldrich, St. Louis, MO, USA) at either 0.1 µg/mL, 1 µg/mL, or 10 µg/mL concentrations. After 48 hrs, cells were treated with 50 µL of 5 mg/mL MTT/1× PBS and incubated for 3 hrs at 37°C in a 5%-CO_2_ incubator. Absorbance was read at 590 nm. All samples were evaluated in triplicate. Five independent analyses were evaluated and mean values were calculated to determine EV size. Data are presented as means ± standard error of means (SEMs).

### 2.6 *In vivo* EV isolation

EV isolation from mouse serum was performed using ExoQuick™ solution (System Biosciences, Palo Alto, CA, USA). Following serum collection, samples were centrifuged at 3000 × g for 15 min to remove cellular debris. Supernatants were transferred to sterile 1.5-mL centrifuge tubes and an appropriate amount of ExoQuick™ solution was added. This mixture was centrifuged at 1500 × g for 30 min at room temperature. Supernatants were then aspirated, and an additional spin at 1500 × g for 10 min was performed. Once residual fluid was removed from each sample, EV pellets were resuspended in 200 μL of sterile 1× PBS. To determine EV protein concentrations, Bradford-Lowry protein quantitation was used (Bio-Rad Laboratories, Hercules, CA, USA).

### 2.7 Nanosight tracking analysis

EV particle size and concentration were measured by Nanosight Tracking Analysis (NTA) (NS3000-LM10, Malvern Instruments, Inc., Malvern, UK) according to the manufacturer’s instructions. In brief, EV samples were diluted by a factor of 1:1000 using filtered sterile 1× PBS. Each sample analysis was conducted for 1 min in triplicate. Five independent analyses were evaluated and mean values were calculated to determine EV size. Data are presented as means ± standard error of means (SEMs).

### 2.8 Enzyme-linked immunosorbent assays (ELISAs)

To detect specific EV proteins, 40 µg of protein samples were used. ELISA plates were coated with either 40 µg of EV samples or blocking buffer (as controls). Protein samples were incubated overnight at 4°C in 96-well plates with bicarbonate buffer (pH 9.5) which bound the vesicles and made the vesicles porous. EV proteins were blocked for 60 min at 4°C in 0.05% bovine serum albumin and Tween 20. After incubation, EV proteins were washed three times and 100 µl of primary antibody was added and incubated for 120 min. ELISA plates were then washed and blocked for 30 min. After blocking, the appropriate secondary antibodies were added. Plates were then washed and signals detected with SIGMAFAST™ *o*-Phenylenediamine dihydrochloride peroxidase substrate (Sigma Aldrich, St. Louis, MO, USA). ELISA plate signals were read at OD 405 nm. Six to eight independent analyses were evaluated and mean values were calculated to determine EV size. Data are presented as means ± standard error of means (SEMs).

### 2.9 Dot blot analysis

Cell lysates were evaluated using 5 µg of lysate, lysate was boiled and blotted on nitrocellulose membranes for 10 min. Samples were blocked in a Pierce Fast-Blocker with 0.09% Tween-20 for 5 min. After blocking, primary CD81 (Thermo Fisher Scientific, Waltham, MA, USA, 1:500) were added to the samples for incubation. After 1 h of incubation at RT, nitrocellulose blots were washed three times with 0.09% Tween-20 in 1× PBS for 10 min. Secondary antibody, Goat anti-rabbit heavy and light chain (H+L) secondary antibody Horseradish peroxidase (HRP) – conjugated rabbit anti-hamster (1:2000 dilution, (Thermo Fisher Scientific, Waltham, MA, USA)) was added in blocking solution (0.09% Tween-20 in 1× PBS) for 1 h and incubated at RT. The blots were washed three times with 0.09% Tween-20 in 1× PBS for 10 min. The nitrocellulose membranes were developed using SuperSignal West Femto Maximum Sensitivity Substrate (Thermo Scientific, Waltham, MA, USA). The signals were read on a Bio-Rad ChemiDoc XRS+ System (BioRad Laboratories, Hercules, CA, USA) using chemiluminescence.

### 2.10 Animals

Eight female BALB/c mice, 6–8 weeks old, were purchased from Charles River Laboratories and maintained in a dedicated animal facility, in sterile cages, with food and water *ad libitum*. All animal experiments were performed in accordance with approved animal regulations and guidelines established by the Institutional Animal Care and Use Committee (Protocol #053116).

### 2.12 *In vivo* inoculation

A low-virulence experimental model was used. Mice were inoculated by intraperitoneal injection of 1 mg/mL bacterial LPS. Control mice were inoculated only with sterile 1× PBS. Final inoculation volumes were adjusted to 400 μL with sterile PBS. Ten days after inoculation, blood (500 µL) was collected via retro-orbital bleeding with anesthetization. Following blood collection, animals were euthanized.

### 2.12 Statistical analyses

All data were analyzed with GraphPad Prism 5 (San Diego, CA, USA) software using a one-way ANOVA with Tukey post-hoc analyses. The Tukey multiple-comparison tests were performed with significance set at *p < 0.05, **p < 0.01, and ***p < 0.0001. Data are displayed as mean ± standard deviation or standard error (as indicated).

## 3. Results

### 3.1 Bacterial LPS alters cell viability

The first set of experiments was designed to investigate the effect of exposing A549 cells to bacterial LPS. Using the MTT assay, cells were treated with different concentrations (0.1 µg/mL, 1 µg/mL, and 10 µg/mL) of bacterial LPS for 48 hrs. This range of LPS concentrations was chosen to elicit an *in vitro* immune response while evaluating exosomal composition. As shown in Figure 1, cell viability increased after LPS treatment at 0.1 µg/mL and 1 µg/mL. At 0.1 µg/mL concentration there was a significant decrease in viability compared to 10 µg/mL (p≤ 0.05). These results indicate that, at the highest dose of LPS, cell viability was reduced compared with control cells.

**Figure 1.**
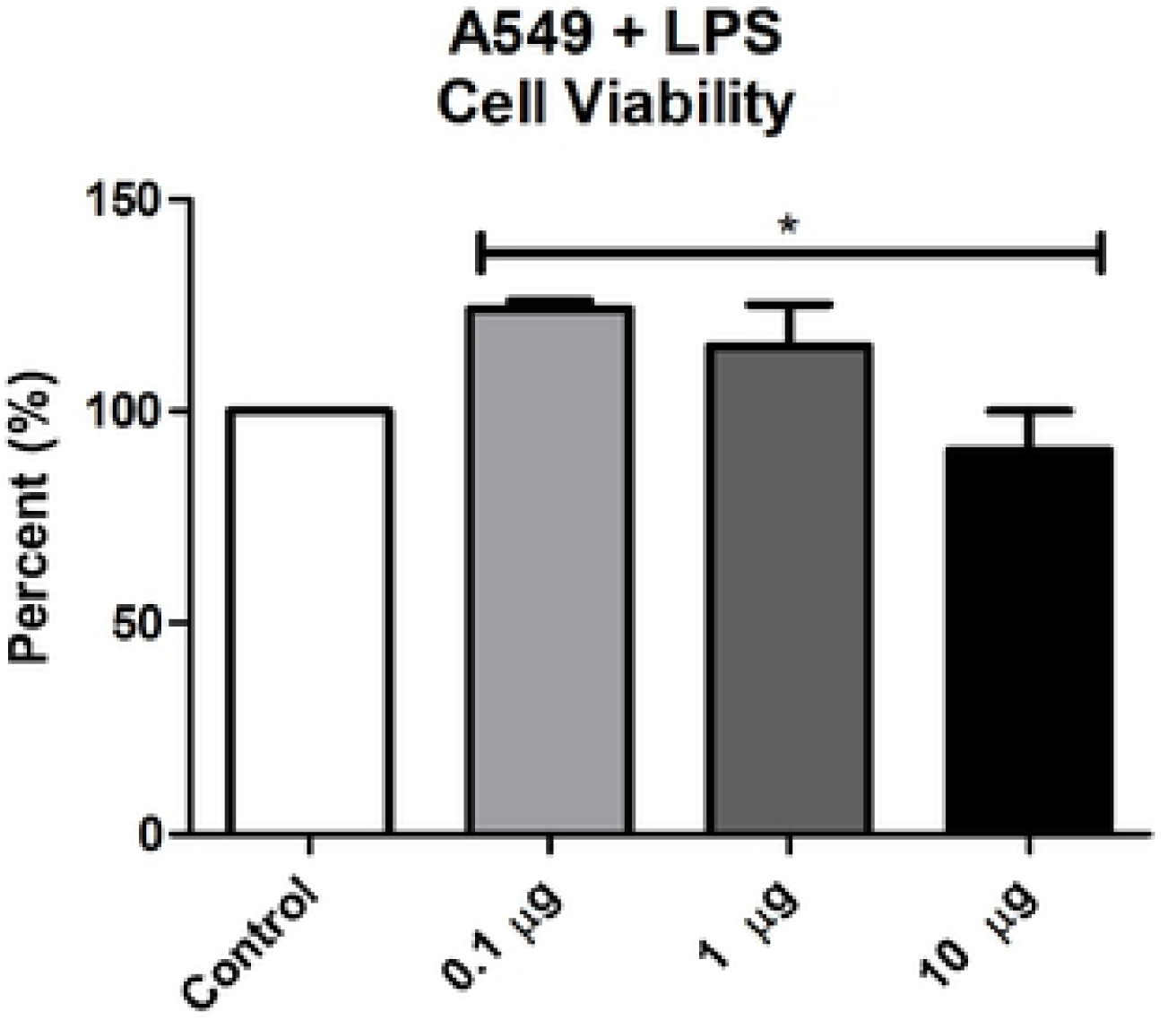
LPS treatment alters cell viability. Cell viability following LPS treatment (0.1 µg/mL, 1 µg/mL, and 10 µg/mL) or without treatment was determined using the MTT assay at 48 h. Mean ± SEM data are from five experiments.

### 3.2 Characterization of EVs from LPS-treated A549 cells

EVs isolated from LPS-treated cells were identified via NTA as vesicles ranging in size from 81–142 nm; 81.3 nm (control), 133.26 nm (0.1 µg/mL), 126.067 (1 µg/mL), and 142.983 (10 µg/mL) (Figure 2A). The mean size of LPS-treated cell derived EVs were significantly increased at 0.1 µg/mL (p ≤ 0.05) and 1 µg/mL (p ≤ 0.01) LPS inoculation when compared to the control-derived EVS. After 48 hrs, the concentration of control A549-derived EVs was 3.48 × 10^8^ particles/mL (Figure 2B). At the lowest concentration of LPS (0.1 µg/mL) there was a significant decrease in particle concentration (6.5 × 10^7^ particles/mL; p ≤ 0.05) compared with control cells (Figure 2B). After 1 µg/mL LPS administration, particle concentration significantly decreased (p ≤ 0.05) to 1.15 × 10^8^ particles/mL and then decreased to a concentration of 1.96 × 10^8^ particles/mL with 10 µg/mL LPS treatment when compared to the control-derived EVs (Figure 2B). These findings suggest that LPS exposure greatly reduced the number of EVs being released from cells in response to pathogenic virulence factors.

**Figure 2.**
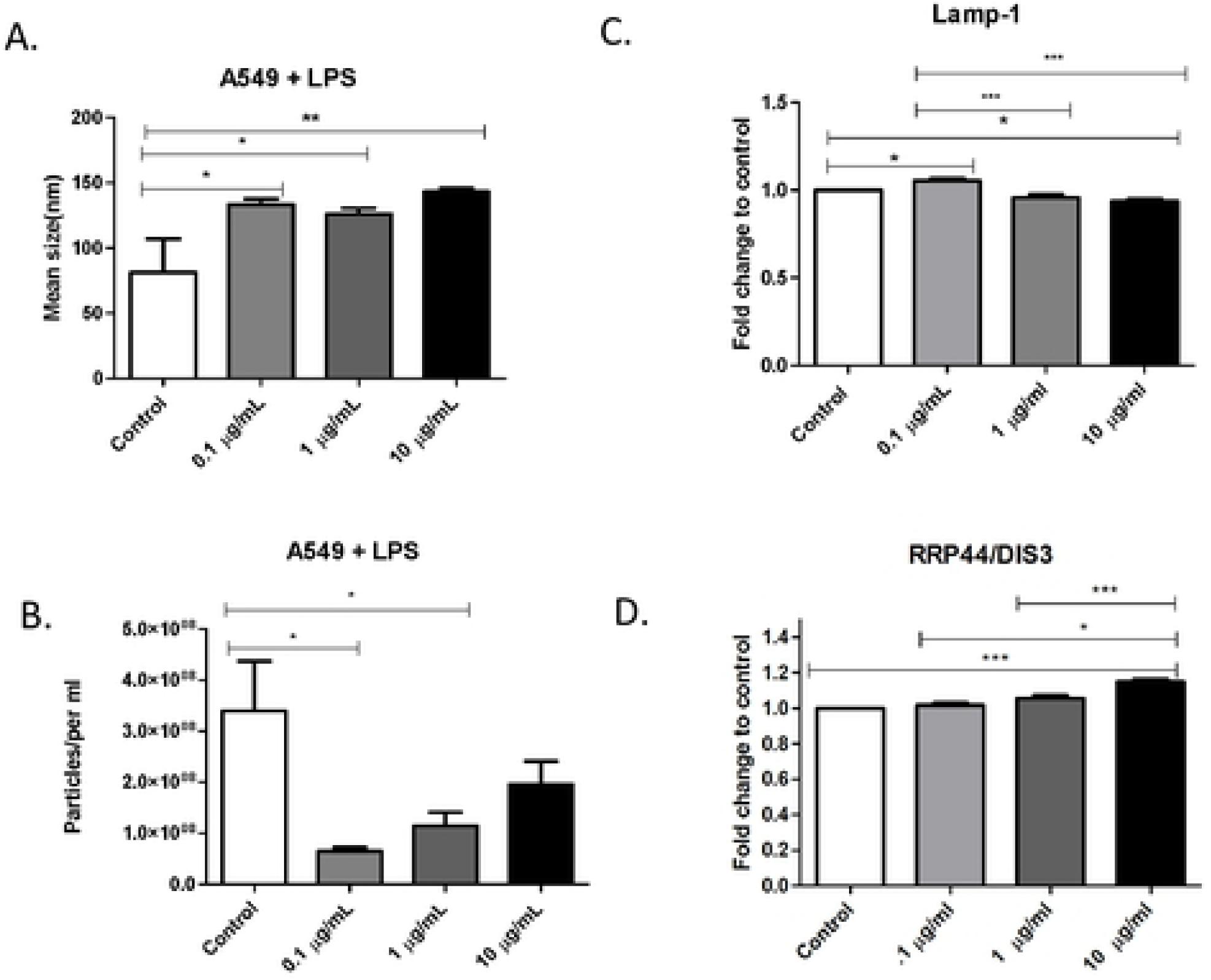
LPS treatment alters EVs from A549 cells. (A) Mean sizes and (B) particle concentrations (per mL) were determined for A549-derived EVs following LPS treatment using Nanosight Tracking Analysis. ELISAs of A549-derived EVs were used to examine expressions of (C) LAMP-1, and (D) RRP44/DIS3 proteins. Mean fold change ± SEM data are from a total of five experiments. *p ≤ 0.05, **p ≤ 0.01, and ***p ≤ 0.001.

Vesicles from A549-derived EVs expressed the EV marker, LAMP-1 (Figure 2C). LAMP-1 expression was significantly expressed from control to 0.1 µg/mL LPS administration (p ≤ 0.05). The level of EV LAMP-1 from 0.1 µg/mL-treated cells decreased significantly (p ≤ 0.001) compared with that of 1 µg/mL-treated cells and significantly decreased (p ≤ 0.001) from 0.1 µg/mL to 10 µg/mL. 10 µg/mL LPS-derived EVs were also significantly decreased (p ≤ 0.05) compared to the control. Exosomal complex exonuclease RRP44/DIS3 was also expressed in A549 cell-derived EVs (Figure 2D). Compared to control-cell EV levels, RRP44/DIS3 increased significantly (p ≤ 0.001) after 10 µg/mL LPS treatment (Figure 2D). In addition, the presence CD81, an EV-associated protein, was observed in the cell lysate via dot blot analysis, confirming the successful collection of EVs (Supplemental Figure 1). These results demonstrate the successful harvest of EVs derived from treated A549 cells.

### 3.3 Proinflammatory responses from LPS-stimulated A549 cells

LPS recognition by the immune system is a fundamental step for recognizing invading pathogens and for the initiation of an immune response. Anti-inflammatory cytokines are immunoregulatory molecules that control the proinflammatory cytokine response and play a major role in physiological systems, including the nervous system [12, 13]. Cytokines act in conjunction with specific cytokine inhibitors, such as TNFα and soluble cytokine receptors, to regulate the human immune response [13]. LPS elicits the synthesis of inflammatory mediators, which are known to regulate the acute-phase response [14]. In order to investigate cytokine expression in EVs from A549 cells, we used ELISAs to demonstrate the presence of IL-6, IL-1β, TLR4, TNFα, and iNOS (Figure 3A–E).

**Figure 3.**
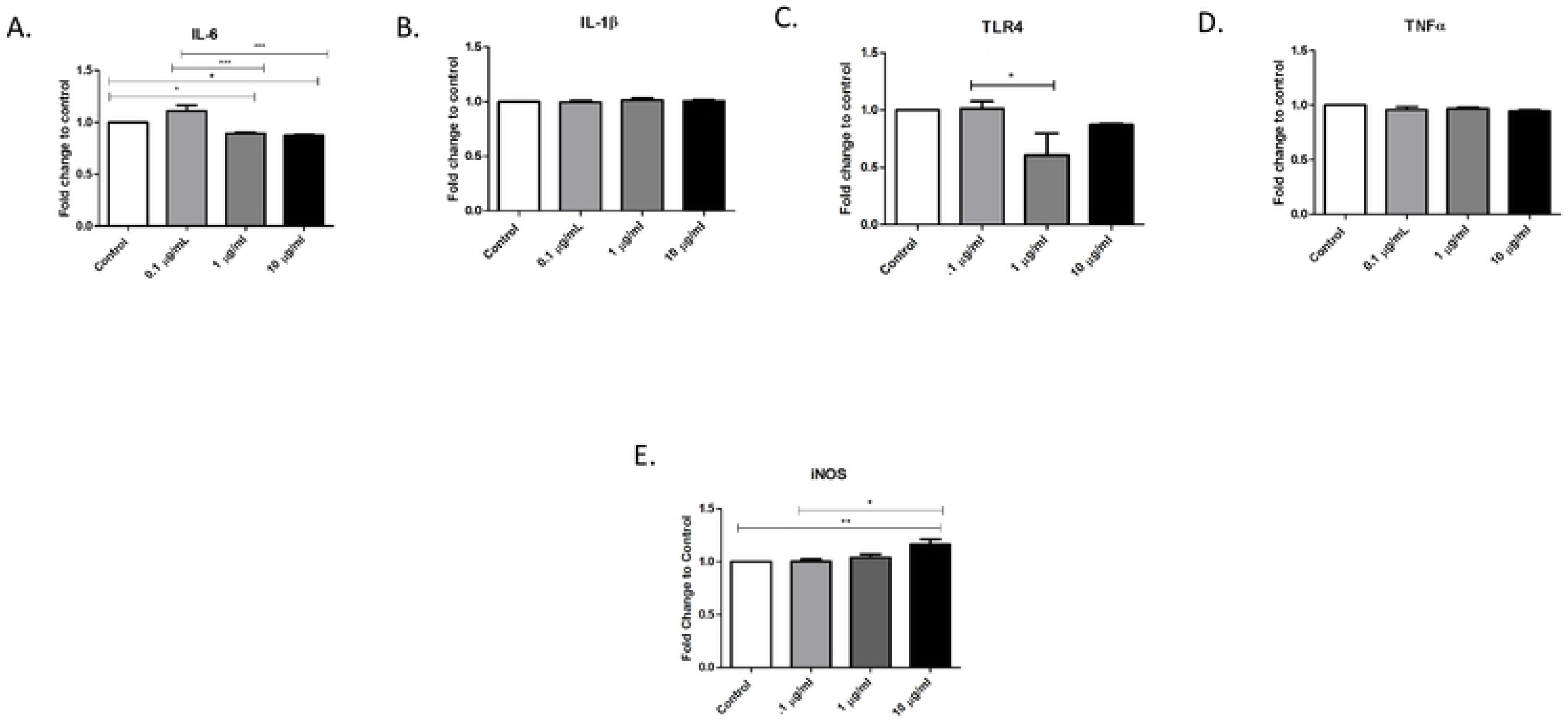
Immunomodulator responses in EVs from LPS-treated cells. The expressions of immunomodulators (A) IL-6, (B) IL-1β, (C) TLR4, (D) TNFα, and (E) iNOS in EVs from LPS-treated (0.1 µg/mL, 1 µg/mL, and 10 µg/mL) A549 cells and untreated cells were determined by ELISA. Mean fold change ± SEM data are from a total of 6–8 experiments. *p ≤ 0.05, **p ≤ 0.01, and ***p ≤ 0.001.

In A549 cell-derived EVs, IL-6 was found to be significantly decreased (p ≤ 0.05) in both 1 µg/mL and 10 µg/mL LPS-treated cells compared with EVs from control cells. Similarly, the level of IL-6 in EVs from 0.1 µg/mL LPS cell-derived EVs was also decreased significantly (p ≤ 0.001) in both 1 µg/mL and 10 µg/mL LPS cell-derived EVs (Figure 3A) when compared with that in EVs from control cells. There was no difference in IL-1β expression between EVs from control cells and EVs from LPS-treated cells (Figure 3B). TLR4 induction has been found in epithelial cells (including lung cells) that are in contact with the external environment [15], and pathogenic infections could potentially affect lung cell physiology through TLR4-mediated local infection or inflammatory responses [16]. Compared with control cell-derived EVs, EVs from cells treated with 1 µg/mL LPS showed a significant decrease (p ≤ 0.05) in TLR4 expression (Figure 3C). The early response cytokine, TNFα, also showed expression in EVs from LPS-treated cells (Figure 3D). These data support the idea that EVs act as carriers for cytokines in response to infection [15]. Compared to control cell-derived EVs, EVs from 10 µg/mL-treated cells showed a significant increase in the expression of the host defense mediator, iNOS (Figure 3E).

### 3.4 Characterization of *in vivo* EVs after LPS treatment

We evaluated the effects of LPS treatment on EV composition collected from BALB/c mice. Mice were injected via intraperitoneal injection with bacterial LPS (1 mg/mL) or PBS (control) the bacterial LPS was diffused into surrounding tissues including the lungs and carried throughout the blood similarly to gram-negative bacteria’s hematogenous mechanism. Ten days later, their blood was collected by retro-orbital bleed. EVs were isolated from serum using the ExoQuick™ process, and *in vivo* EV production was evaluated by NTA. The particle analysis data showed that average control EV size was 125 ± 0.9 nm and LPS-derived EV size was 115 ± 5.04 nm (Figure 4A). Compared to the EV concentration in control mice (5.18 × 10^8^ ± 1.9 × 10^7^ particles/mL), there was a slight decrease (Figure 4B) in EV concentration (4.54 × 10^8^ ± 8.7 × 10^6^ particles/mL) in LPS-treated mice. These data indicate a slight decrease in EV size and particle concentration in the LPS-treated group compared to the control group, suggesting that exposure to LPS slightly decreased the number and mean size of EVs being released.

**Figure 4.**
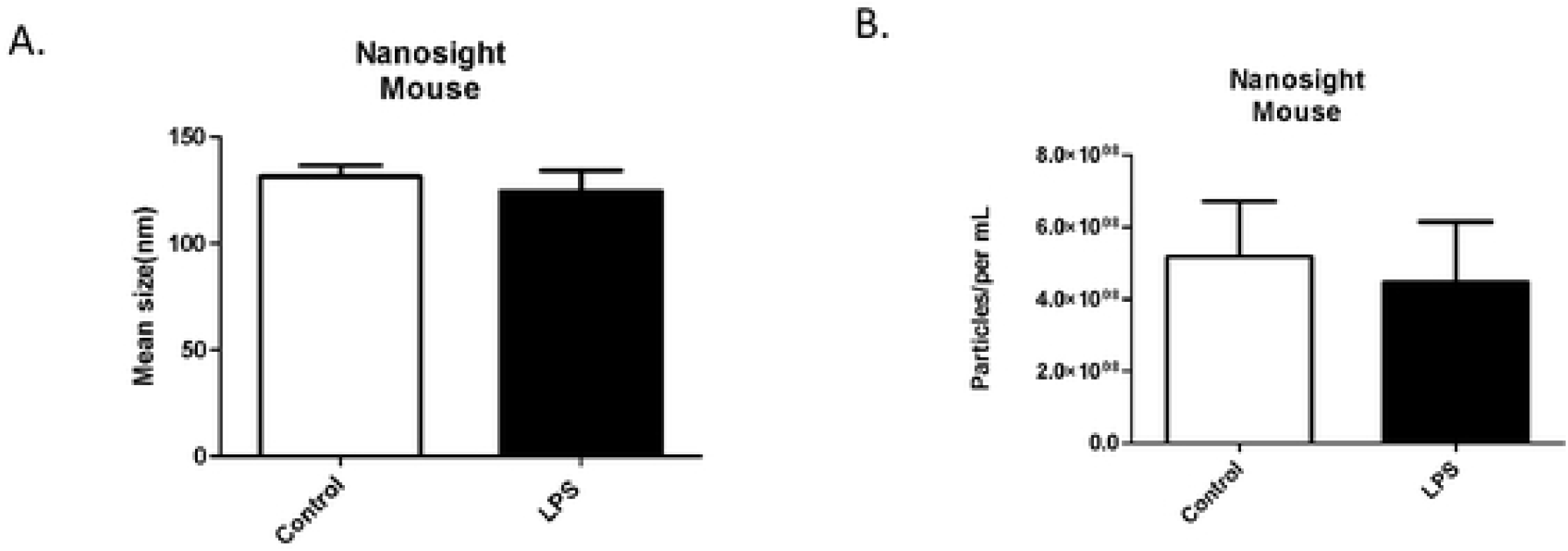
EV characterization from LPS-treated mice *in vivo*. (A) Mean sizes and (B) particle concentrations (per mL) were determined for mouse-derived exosomes following LPS treatment using Nanosight Tracking Analysis. Mean ± SEM data are from a total of eight control mice and six experimental mice.

### 3.5 EV-associated proteins *in vivo*

Members of the tetraspanin superfamily play fundamental roles in a multitude of biological processes including cell motility, invasion, adhesion, and protein trafficking [17]. EVs are highly enriched in tetraspanins and they are frequently used as EV markers, so we examined EV-associated protein expression after LPS treatment *in vivo*. EVs from the LPS-treated group expressed CD9 (Figure 5A), CD81 (Figure 5B), and CD63 (Figure 5C). CD81, frequently found in EVs, was significantly (p ≤ 0.01) increased in the LPS-treated group compared with the control group (Figure 5B). The expression of the EV-associated protein, LAMP-1, was also increased in LPS-treated EVs (Figure 5D). These data indicate that EV markers are shuttled into EVs in response to LPS treatment, further confirming the successful isolation of EVs.

**Figure 5.**
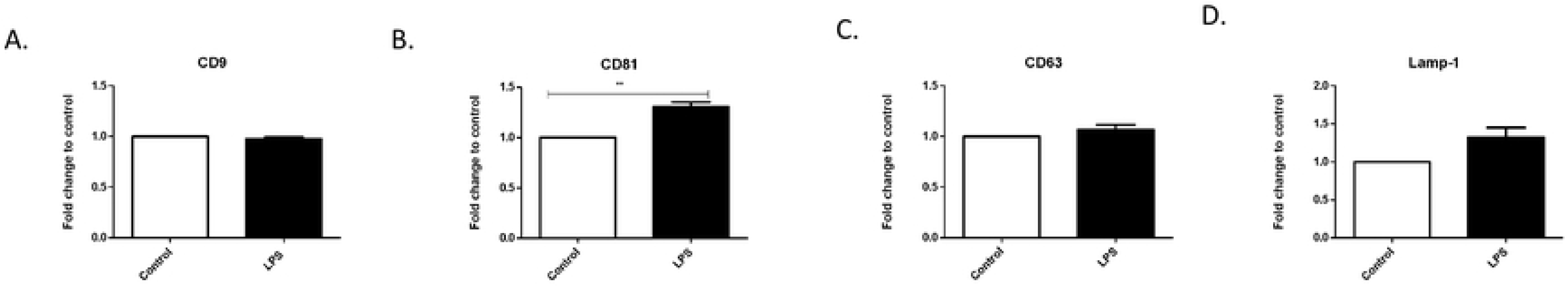
EV-associated protein expression from LPS-treated BALB/c mice. The expressions of (A) CD9, (B) CD81, (C) CD63, and (D) LAMP-1 were determined by ELISA in EVs isolated from mice. Mean fold change ± SEM data from a total of five experiments. **p ≤ 0.01.

### 3.6 Immunomodulator response to LPS *in vivo*

Systemic exposure to an endotoxin such as LPS results in the release of active inflammatory mediators, such as IL-6, through cell-signaling pathway cascades. Here, we found the presence of IL-6 in both control EVs and those isolated from LPS-treated mice (Figure 6A). There was no significant difference between untreated and LPS-treated mice. The host immune response system includes TLRs that can recognize LPS. Specifically, TLR4 is the main receptor responsible for initiating a response to LPS in the outer membrane of gram-negative bacteria. Thus, we sought to evaluate the expression of these important receptors in EVs derived from LPS-treated mice. Both TLR4 and TLR7, innate immune response proteins, were observed, and TLR4 expression was significantly decreased when compared to the control (p ≤ 0.001) (Figure 6B and C). These data suggest that EVs may act as transport vehicles for cytokines and chemoattractants for natural killer cells.

**Figure 6.**
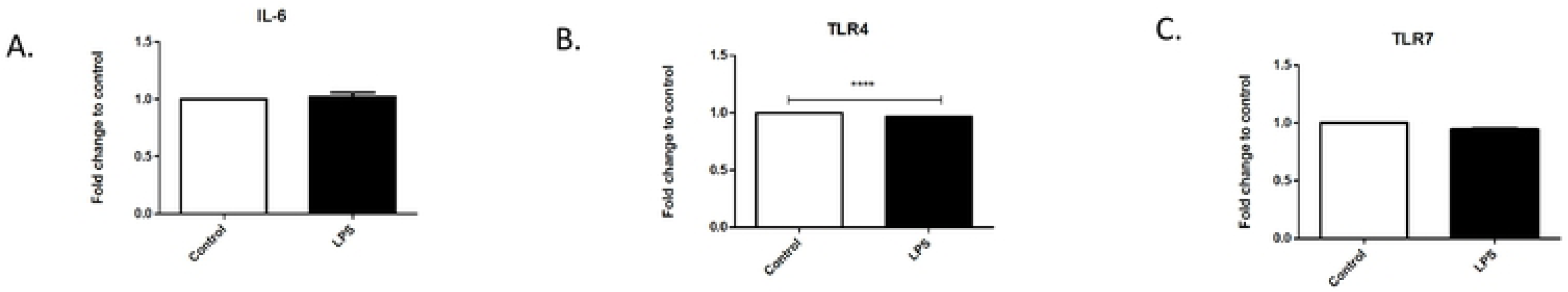
Expression of immunomodulators after LPS treatment *in vivo*. The expressions of (A) IL-6, (B) TLR4, and (C) TLR7 were determined by ELISA in EVs isolated from mice. Mean fold change ± SEM data from a total of five experiments. ****p ≤ 0.0001.

## 4. Discussion

Gram-negative bacteria represent a major group of pathogens causing numerous human diseases (e.g., pneumonia and meningitis), and the LPS endotoxin is an important mediator of septic shock, a major cause of death in critical-care facilities worldwide. Sepsis involves an uncontrolled inflammatory response by host cells that can lead to organ failure and subsequent death [18]. As of 2016, septic shock prevalence was estimated to be 300 cases per 100,000 people in the United States, resulting in 200,000 deaths annually [19]. Gram-negative pathogens can evade immune defenses and spread to other organs by producing an extensive array of virulence factors (e.g., LPS). These virulence factors interact with host cells by using specific receptors to interfere with the host immune response. Understanding the role of LPS in pathogenic invasion and the host immune response has become increasingly important for therapeutic treatment. Here we examined the role of EVs in disease progression by evaluating the effect of bacterial LPS on immune responses *in vitro* and *in vivo*. The role of EVs released by stimulated lung cells has not been well studied, so we first assessed the effects of LPS treatment on EVs derived from A549 lung cells and those circulating in mouse serum.

We first examined EV size and particle number after treatment with three different concentrations of bacterial LPS (0.1 µg/mL, 1 µg/mL, and 10 µg/mL) using cultured lung cells. For EVs derived from these cells, their mean size at all concentrations of LPS treatment showed a significant increase compared with that of control EVs from untreated cells. This size increase suggests an addition of EV cargo by LPS stimulation. The concentration of EVs from untreated cells was 3.4 × 10^8^ particles/mL and decreased significantly (p ≤ 0.05) after 0.1 µg/mL LPS treatment. Particle concentrations then increased in the 1 µg/mL and 10 µg/mL-treated cells compared with the lowest level LPS treatment. These increases in EV concentration after LPS treatment differed from a recent report examining EVs after LPS treatment of cardiomyocytes [23]. In that study, LPS treatment caused cardiomyocytes to release EVs at a slower rate compared to their controls, suggesting that insult-induced EV release may be cell-type specific and be dependent on regional anatomy [11].

We also characterized EV proteins from control and inflammatory environments *in vitro* and *in vivo*. Using ELISAs, we observed changes in the presence of EV markers, immunomodulators, and stress-associated proteins (data not shown) to elucidate the role these vesicles might play in disease progression [20]. The exosome-associated marker, LAMP-1, was significantly expressed in A549-derived EVs after LPS treatment. RRP44/DIS3 was also significantly expressed in EVs from these cells. Combined with the significant increase in EV size after LPS treatment observed using NTA, this suggests that EVs released under inflammatory conditions have increased protein packaging to aid the host’s immune system. We further confirmed the presence of EVs by examining the expression of RRP44/DIS3 in EVs from treated and untreated cells. RRP44/DIS3 was significantly increased (p ≤ 0.001) in EVs from 10 µg/mL-treated cells compared with EVs from control cells. RRP44 is found in the extracellular vesicle complex in the nucleus of eukaryotic cells and plays a role in the recognition of, as well as degradation of, RNA targets.

The respiratory airway is constantly exposed to pathogens and LPS. The body must react immediately to these pathogens to prevent subsequent infections such as pneumonia. Bacterial LPS exposure might also induce an excessive inflammatory response that could ultimately be detrimental to the host. Interleukins play a vital role in host protection by activating adaptive immunity at the beginning of pathway signaling for apoptosis. IL-6, a multifunctional cytokine, is present in EVs from A549 cells. Interleukins are expressed in a variety of tissues and are known to exert their influence through the circulatory system [21]. In A549 cell-derived EVs, IL-6 expression was slightly increased compared with control expression after 0.1 µg/mL LPS treatment, but was significantly reduced (p ≤ 0.05) after 1 µg/mL and 10 µg/mL LPS treatments compared with control EVs. There were also significant IL-6 differences (p ≤ 0.001) between EVs from 1 µg/mL LPS-treated cells and from cells treated with the other two LPS concentrations. IL-6 was also found in EVs isolated from serum *in vivo*. It is possible that increasing the number of LPS injections in these mice might have further increased IL-6 expression to levels reported by Erickson et al. [29], where significant IL-6 expression was seen when mice were given three LPS injections [22]. The pro-inflammatory cytokine IL-1β induces many inflammatory pathways, including the stimulation of other cytokines and chemokines, recruits neutrophils to sites of inflammation, and initiates adaptive immunity [23]. IL-1β release has also been shown to be associated with infectious bacteria such as gram-negative *H. influenzae* [24]. While there were no significant treatment differences between the control and LPS treatment groups, IL-1β was present in EVs from A549 cells. TLRs play a vital role in early innate-immunity responses to invading microbes by sensing the invasion. In particular, TLR4 stimulation leads to intracellular signaling and inflammatory cytokine activation that triggers host innate immunity. TLR4 expression was seen in EVs from A549 cells and decreased slightly in EVs from 1 µg/mL-treated cells, but expression increased in EVs from 10 µg/mL-treated cells. Both TLR4 and TLR7 were found in EVs from LPS-treated mice, and TLR4 expression was significantly reduced (p ≤ 0.0001) compared with that in EVs from control mice. iNOS expression increased significantly (p ≤ 0.001) in response to LPS treatment. TNFα was also present in A549 cell-derived EVs. These results indicate that immunomodulators are present in EVs after LPS treatment and suggest that cellular responses to pathogenic infections initiate these changes to EV cargo. The presence of inflammatory agents and inflammatory response stimulators in EVs derived from LPS-stimulated cells suggests that they are key EV molecules that respond to pathogenic invasions.

*In vivo*, we treated BALB/c mice with 1 mg/mL bacterial LPS and examined serum-derived EVs for EV-associated markers. We found that tetraspanins, CD9, and CD63 were all represented in these EVs, and CD81 (p ≤ 0.05) was significantly so. Tetraspanins have the capacity to form multifunctional tetraspanin-enriched microdomains that are essential mediators for EV interactions, selection of vesicle cargo, antigen presentation during immune responses, and binding and uptake of EVs by target cells [25]. As further confirmation of successful EV isolation, LAMP-1 protein expression was higher in EVs from LPS-treated mice than in EVs from untreated mice. As LAMP-1 expression is a marker for late endosomes, this suggests that EVs have undergone endocytosis and been transported to endosomal organelles. In our mouse model, the mean size of our control EVs was 131 nm. After LPS treatment, there was a slight decrease of 7 nm in EV size. Mean particle concentration also decreased from 5.18 × 10^8^ in the control group to 4.4 × 10^8^ particles/mL in the LPS-treated group. This suggests that EVs are being released at different rates by different cell types in response to LPS stimulation.

## Conclusion

EVs provide a means both for initiating an immune response and for being vehicles for agents that promote disease progression. They can play a key role in intracellular communication and for influencing the phenotype of recipient cells. EV studies are still in their infancy, but progress in this field has provided new prospects for understanding how activation of the innate immune system causes life-threatening human infections. In addition, a new chapter has been opened with the discovery of a link between EVs and LPS-induced stimulation, but further studies on this topic are needed. The most noteworthy future application of EVs will be to harness and manipulate their cargo as therapeutic tools to monitor and diagnosis disease. Considerable research will be needed to accomplish this goal.

## Author contributions

Formal analysis, Q.L.M.; Funding acquisition, Q.L.M.; Investigation, C.R.B., L.B.J., B.J.C., S.K., and Q.L.M.; Methodology, C.R.B., L.B.J., B.J.C., S.K., M.T.C, and Q.L.M.; Supervision, Q.L.M.; Writing (original draft), C.R.B., L.B.J., and B.J.C.; Writing (review and editing), C.R.B and Q.L.M.

## Funding

This work was funded by the National Institutes of Health, #1R15DA045564-01 (QLM) and National Science Foundation’s Alliances for Graduate Education and the Professoriate (AGEP) Program (LBJ), National Science Foundation #Grant Nos. IOS-1432991 (QLM), National Science Foundation-HBCU-RISE (HRD-1646729) (LBJ).

### Acknowledgements

Special recognition to the High-Resolution Imaging Facility Service Center for providing the NTA at the University of Alabama at Birmingham, and the following grants: Cancer Center Support Grant P30 CA013148; Rheumatic Disease Core Center P30 AR048311.

## Data Accessibility

The original data will be maintained by the corresponding author. Information pertaining to the data sets will be made available upon written request.

---

**Supplemental Figure 1.**
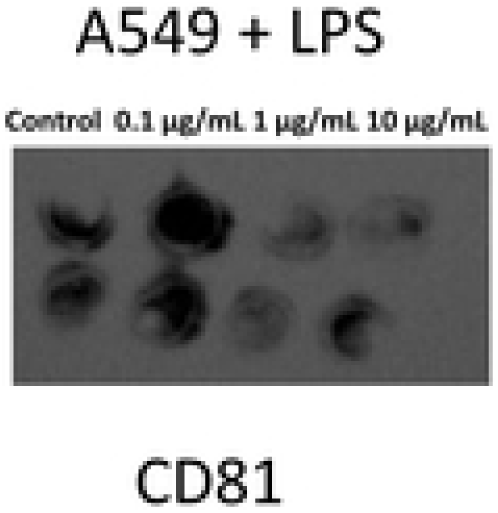
Expression of an EV-associated protein in A549 cell lysates. Dot blot analyses of CD81 in A549 cell lysates following LPS treatment (0.1 µg/mL, 1 µg/mL, and 10 µg/mL).

